# Structure-conditioned masked language models for protein sequence design generalize beyond the native sequence space

**DOI:** 10.1101/2023.12.15.571823

**Authors:** Deniz Akpinaroglu, Kosuke Seki, Amy Guo, Eleanor Zhu, Mark J. S. Kelly, Tanja Kortemme

**Affiliations:** The UC Berkeley-UCSF Graduate Program in Bioengineering, University of California, San Francisco, CA; Department of Bioengineering and Therapeutic Sciences, University of California, San Francisco, CA; Department of Pharmaceutical Chemistry, University of California, San Francisco, CA; Quantitative Biosciences Institute, University of California, San Francisco, CA; Chan Zuckerberg Biohub, San Francisco, CA

**Keywords:** deep learning, masked language modeling, protein design, novel sequence design

## Abstract

Machine learning has revolutionized computational protein design, enabling significant progress in protein backbone generation and sequence design. Here, we introduce Frame2seq, a structure-conditioned masked language model for protein sequence design. Frame2seq generates sequences in a single pass, achieves 49.1% sequence recovery on the CATH 4.2 test dataset, and accurately estimates the error in its own predictions, outperforming the autoregressive ProteinMPNN model with over six times faster inference. To probe the ability of Frame2seq to generate novel designs beyond the native-like sequence space it was trained on, we experimentally test 26 Frame2seq designs for de novo backbones with low identity to the starting sequences. We show that Frame2seq successfully designs soluble (22/26), monomeric, folded, and stable proteins (17/26), including a design with 0% sequence identity to native. The speed and accuracy of Frame2seq will accelerate exploration of novel sequence space across diverse design tasks, including challenging applications such as multi-objective optimization.

## 1. Introduction

The ability to design proteins with new functions has widespread applications in biotechnology and medicine. Traditionally, computational protein design has relied on physics-based principles and simulations. Recently, machine learning methods learning directly from data have enabled promising advances, such as highly accurate structure prediction with AlphaFold2 (1) and comparable methods (2–4), as well as inverting these models for protein design (5–10).

Computational protein design has achieved high experimental success rates in diverse applications, including generation of new protein folds (5, 9) and symmetrical oligomers (9, 11), protein-protein binder design (9), and motif-scaffolding (6, 9). Essentially all of these methods first generate a protein backbone followed by fixed backbone sequence design as a second step.

ProteinMPNN is a state-of-the-art encoder-decoder model that autoregressively predicts protein sequences given a backbone (12). ProteinMPNN has been experimentally validated to yield successful monomeric proteins and protein assemblies. Alternative methods built on encoder-only architectures have also been proposed, such as PiFold (13). PiFold outperforms ProteinMPNN on native sequence recovery but has not been experimentally validated (13).

In this work, we introduce Frame2seq, a structure-conditioned masked language model that designs protein sequences. Frame2seq is an invariant point attention (IPA)-based bidirectional encoder and achieves fast inference due to single-pass sequence generation. Through in silico and experimental evaluation, we demonstrate that Frame2seq is a state-of-the-art fixed-backbone design method and produces soluble, monomeric, and stable proteins. Importantly, we demonstrate that our model pseudo-log-likelihood is highly correlated with prediction success. We also test and experimentally validate the ability of Frame2seq to design novel sequences with undetectable similarity to the starting protein, showing that Frame2seq generalizes beyond the sequence space it was trained on. This ability to generate sequence novelty has fundamental implications for questions on the size of the designable sequence space and enables particularly challenging applications of sequence design such as multi-objective optimization.

## 2. Methods

### Datasets

We train Frame2seq on the non-redundant CATH 4.2 (14) data splits of single chain proteins up to length 500 as described by (15). There are 18024, 608, and 1120 chains with no topological overlap in the training, validation, and test sets, respectively. These splits are clustered for structural diversity and allow to test generalization across many folds (15). To perform direct comparisons, we benchmark against models trained on the same dataset.

### Structure-conditioned masked language model

Frame2seq is a translation- and rotation-invariant encoder-only model that takes protein backbones as input and generates protein sequences (Figure 1A). In a single inference pass, Frame2seq designs complete protein sequences. We preserve the invariance property using IPA (1). We compute rotations and translations from input backbone coordinates to construct coordinate frames as described for AlphaFold2 (1). To obtain sequence embeddings, we compute phi, psi, and omega torsions, along with absolute position embedding. To obtain pairwise embeddings, we compute inter-residue distances lifted into a radial basis and relative position indices between pairs of residues. Coordinate frames, sequence embeddings, and pairwise embeddings pass through a repeating stack of node and edge update operations followed by a final node update and transition to sequence dimension. IPA layers and transition layers comprise the node updates. We update edges by passing pairwise embeddings and updated sequence embeddings into two layers of MLP. Frame2seq is trained to minimize a categorical cross entropy loss for a native sequence recovery objective.

**Fig. 1.**
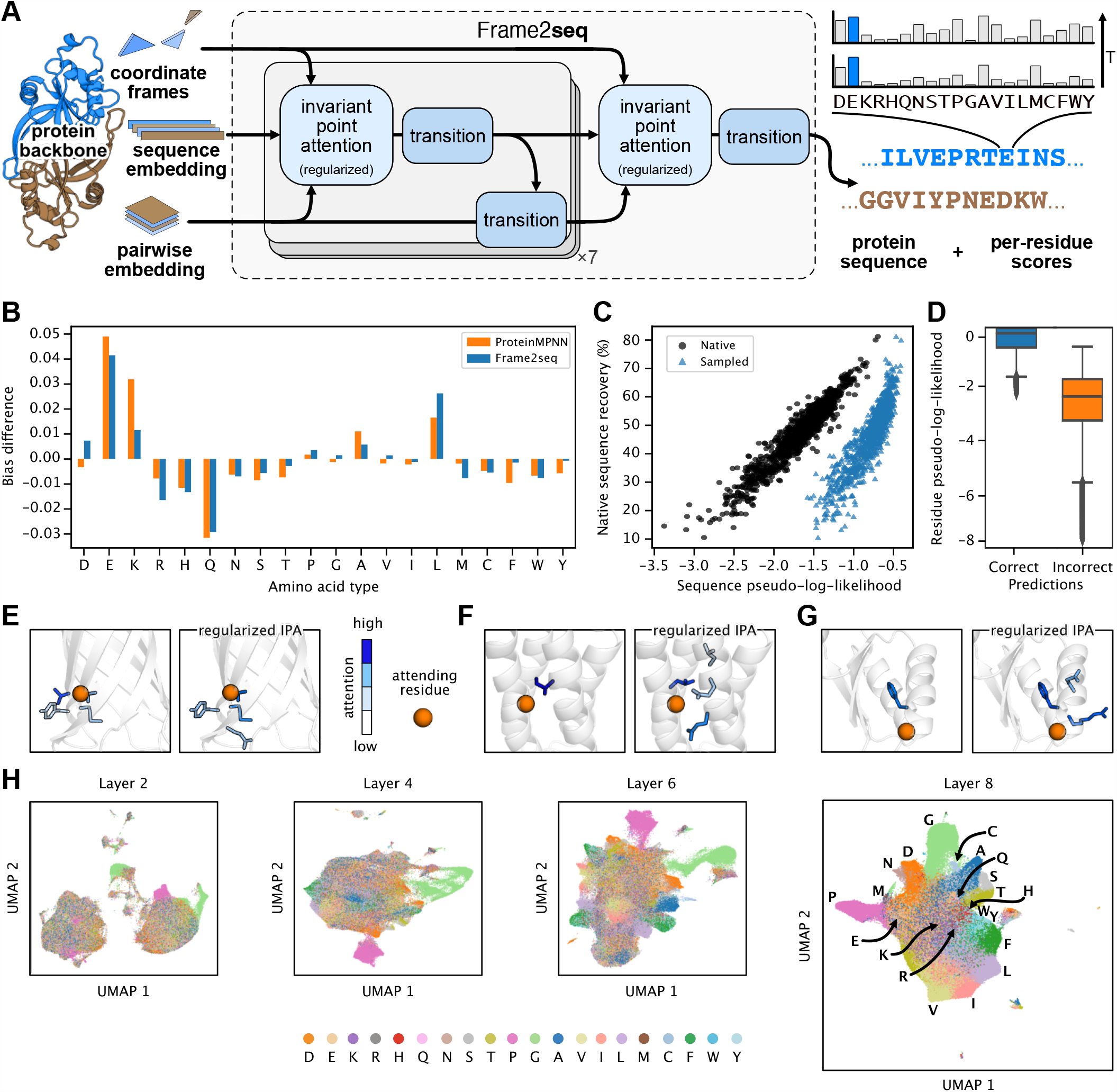
(A) Model architecture. Frame2seq is composed of regularized invariant point attention layers and sequence embedding transitions followed by pairwise embedding updates. Arrows indicate information flow through model components. (B-H) In silico analysis of Frame2seq. (B) Difference in compositional bias between ProteinMPNN and Frame2seq. (C) Correlation between Frame2seq native sequence recovery and model pseudo-log-likelihood averaged over sequence for native (dark blue, Spearman correlation = 0.94) and sampled (light blue, Spearman correlation = 0.89) sequences. (D) Per-residue model negative pseudo-log-likelihood for correct residue identity (native) and incorrect residue identity (non-native) predictions. Incorrect prediction outliers below 3 times the interquartile range are not shown. (E-G) Frame2seq attends to second-shell interactions, bulky side chains, and side chains pointing towards attending residue with IPA regularization (right). Distribution of attention to residues by the attending residue (orange sphere) with IPA (left) and regularized IPA (right). Attention value increases from white to dark blue. Side chains are only shown for residues with attention. (E) PDBID 6X1K. (F) PDBID 1P68. (G) PDBID 2LV8. (H) UMAP representation of Frame2seq embeddings from layers 2, 4, 6, and 8 for test set targets (colored by predicted amino acid identity).

### Attention regularization for IPA layers

We hypothesize that introducing tolerable difficulty during training allows for better model performance. To increase training difficulty without halting the model’s ability to minimize the categorical cross entropy loss, we explore feature dropouts and attention regularization for IPA. We implement regularized IPA layers by randomly masking attention weights between pairs of residues during training (Algorithm 1). We find that a masking rate of 20% is optimal and results in improved model performance (Table S2). We attribute this improvement to rendering the training task sufficiently but not overly more difficult.

## 3. Results

### Frame2seq recovers native sequence from structure

We benchmarked Frame2seq against a ProteinMPNN model trained on the CATH 4.2 dataset without backbone noise. We evaluated the performance of both models when provided no ground truth sequence as context and found that Frame2seq was more accurate than ProteinMPNN. Specifically, Frame2seq achieves 49.1% native sequence recovery, outperforming ProteinMPNN by 2.0%. We predicted structures for the native sequences, ProteinMPNN designs, and Frame2seq designs using all 5 AlphaFold2 models with 3 recycles and no templates. We calculated AlphaFold2 high confidence rate (%) and AlphaFold2 success rate (%), both with and without multiple sequence alignments (MSAs). We found that ProteinMPNN and Frame2seq achieve similar high confidence and success rates. Both models are outperformed by the native sequences when MSAs are provided, which is expected given the inclusion of native sequences in the AlphaFold2 training set (Figure S3). When averaged over the test dataset targets, Frame2seq inference is approximately 6.2 times faster than ProteinMPNN (Table 1). This difference is due to Frame2seq’s single-pass and ProteinMPNN’s autoregressive sampling formulation.

**Table 1.** **Computational benchmark of model performance and speed over CATH 4.2 held out test dataset targets. Sequence recovery is median of average. AlphaFold2 high confidence rate is percentage of targets that achieve pLDDT > 90. AlphaFold2 success rate is percentage of targets that achieve LDDT-C**α **> 90. Statistically significant differences (with p-value < 0.05) between models are bolded for AlphaFold2 predictions. GPU speed (s) is mean of average**.

|  | Sequence recovery (%) ↑ | AlphaFold2 high confidence rate (%) ↑ |  | AlphaFold2 success rate (%) ↑ |  | GPU speed (s) ↓ |
| --- | --- | --- | --- | --- | --- | --- |
|  |  | With MSA | Without MSA | With MSA | Without MSA |  |
| Native | — | 66.25 | 2.59 | 48.48 | 1.34 | — |
| ProteinMPNN | 47.10 | 53.21 | **4.82** | 39.02 | 3.21 | 2.71 |
| Frame2seq | **49.11** | 53.21 | 4.02 | 38.39 | 3.04 | **0.44** |

Incorrect residue predictions of fixed-backbone design methods reveal their underlying compositional bias for certain amino acid types. Compositional bias difference is calculated by subtracting the predicted number of residue type occurrences from true and dividing by the total number. We calculated compositional bias difference for ProteinMPNN and Frame2seq for all amino acid types and found that Frame2seq has overall less bias (Figure 1B).

We next investigated how the performance of ProteinMPNN and Frame2seq depended on residue burial. To measure burial, we computed the average Cβ distance for 8 closest residues (Å) (lower for core residues, higher for surface residues). We found that residue burial has a similar effect on both models, with the expected behavior that core residues are easier to recover than surface residues. Frame2seq’s 2.0% native sequence recovery improvement over ProteinMPNN is primarily at the surface residues of the targets (Figure S2).

### Frame2seq accurately estimates the error in its own predictions

We investigated the ability of Frame2seq to discriminate accurate from inaccurate predictions. Towards this goal, we analyzed how well model pseudo-log-likelihood (PLL) correlates with native sequence recovery. For both native and sampled sequences, we computed model PLL and found it to strongly correlate with native sequence recovery (Figure 1C). Frame2seq scores native and sampled sequences favorably when the model predicts accurately. Frame2seq prefers its own predictions over the native sequences, which is expected due to model bias and the possibility of finding alternative sequences that fold into the input structure. We additionally found model PLL to discriminate accurately between native and non-native predictions when computed per position (Figure 1D).

### IPA regularization enforces side chain awareness

With IPA regularization, Frame2seq learns node updates from the context of a restricted local environment. Training with restricted attention capacity contributes to model’s improved performance (Table S2). To investigate the differences in attention distribution that lead to this improvement, we analyzed and visualized the attention matrices for three de novo backbones: all-α, all-β, and α-β-alternating. When attention between pairs of residues is randomly dropped during training, the model attends to alternative residues that provide the most context for sequence prediction (Figure 1E-G). We found that this change enforced Frame2seq to allot more attention to residues with bulkier side chains (Figure 1E-G), alternative residues with side chains oriented towards the attending residue (Figure 1F), and longer-range second shell interactions (Figure 1F-G).

### Sequence embeddings map physicochemical properties

We visualized Frame2seq sequence embeddings for each layer in two-dimensional UMAP representations colored by the predicted residue identities (Figure 1H). Glycine and proline residues have distinct sequence embeddings in every layer (Figure S1), which we attribute to flexibility patterned in glycine and restriction patterned in proline backbone coordinates. We observed predictions to begin convergence at layer 4 and saturation of convergence at layer 8. Visualization of layer 8 embeddings suggests strong mapping of residue identities to known physicochemical properties.

### Frame2seq designs stable sequences onto de novo backbones

In silico evaluation of Frame2seq demonstrates its ability to generate realistic protein sequences in principle. However, experimental validation of a design method is essential, because even one or a few incorrect amino acids in a protein, while causing a small difference in sequence recovery, can be detrimental to protein stability. Beyond testing for its ability to output native-like proteins, we tasked our model to design sequences onto de novo backbones to have low (zero to fifty percent) sequence identity to the native sequence. This is a challenging task for a model that only learns from the native sequences associated with ground truth backbones. However, we found that Frame2seq’s generative capabilities extend beyond the naturally occurring sequence space as we successfully designed novel low sequence identity proteins.

### Design and characterization of low sequence identity proteins

We generated low sequence identity sequences for the backbones of the first computationally designed de novo fold, Top7 (PDBID 1QYS) (16) and a de novo Rossmann 2x2 fold (PDBID 2LV8) (17). To achieve low sequence identity to native, we restricted Frame2seq model logits to exclude the true residue identity at each or randomly chosen position(s) before sampling sequences. We then predicted structures for each candidate using all 5 AlphaFold2 models with no MSAs, no templates, and 3 recycles. We filtered our designs by calculating an average pLDDT and computing structural deviation from the true backbones, and selected designs with pLDDT > 89 and predicted backbone heavy atom RMSD < 1.15 Å. We searched our design sequences against non-redundant protein sequences and the PDB database and found minimal to no matches. Frame2seq designs with less than 20% sequence identity to the native yield zero sequence matches, and above 20% identity designs are most significantly matched to the native Top7 or the 2LV8 Rossmann fold, respectively.

We experimentally evaluated twenty-six sequences (eighteen and eight designed onto the Top7 and de novo 2LV8 Rossmann backbones, respectively). Out of the total 26, 25 of our designs express in *E. coli*, 22 are soluble, and 17 are monomeric and folded (Figure 2A). 16 of these experimentally successful designs do not melt at up to 95 °C. We demonstrate experimental success over all sequence identity bins we explored (Figure 2B). Figure 2C-H highlights biophysical characterization of Top7 designs with 0% (“Top0”) and 5% sequence identity and 2LV8 designs with 19% and 37% sequence identity to the starting protein. These designs are monomeric when assessed by size exclusion chromatography (Figure 2C-D), fold into the expected secondary structure as measured by circular dichroism (Figure 2E-F), and do not melt at up to 95 °C apart from the 19% identity 2LV8 design (Figure 2G-H). The 19% identity design melts at 55 °C and refolds reversibly after thermal denaturation. These designs are predicted to maintain a near-identical structure as assessed by AlphaFold2 predictions (Figure 2I-J). This design challenge shows that Frame2seq successfully samples unexplored sequence space, down to 0% sequence identity in the case of Top0. Moreover, the heteronuclear single-quantum coherence (HSQC) spectrum of ^15^N-labelled Top0 revealed that this protein is folded with substantial β-sheet conformation and that the chemical shift dispersion is comparable to Top7 (16) (Figure 3A). The amide proton resonances (between 9 and 10 parts per million) indicate the presence of β-sheet residues. We show that our Top0 design exhibits secondary structure features consistent with Top7 and that Frame2seq can design proteins with 0% sequence identity while likely maintaining a fold identical to native. Finally, we computed a pairwise substitution matrix between Top7 and Top0 residues (Figure 3B) and observed that the substitutions are diverse for each amino acid type with an average blosum62 score difference of *−* 2 (Figure S5). These experimental validations confirm that Frame2seq generalizes to novel proteins beyond the training set.

**Fig. 2.**
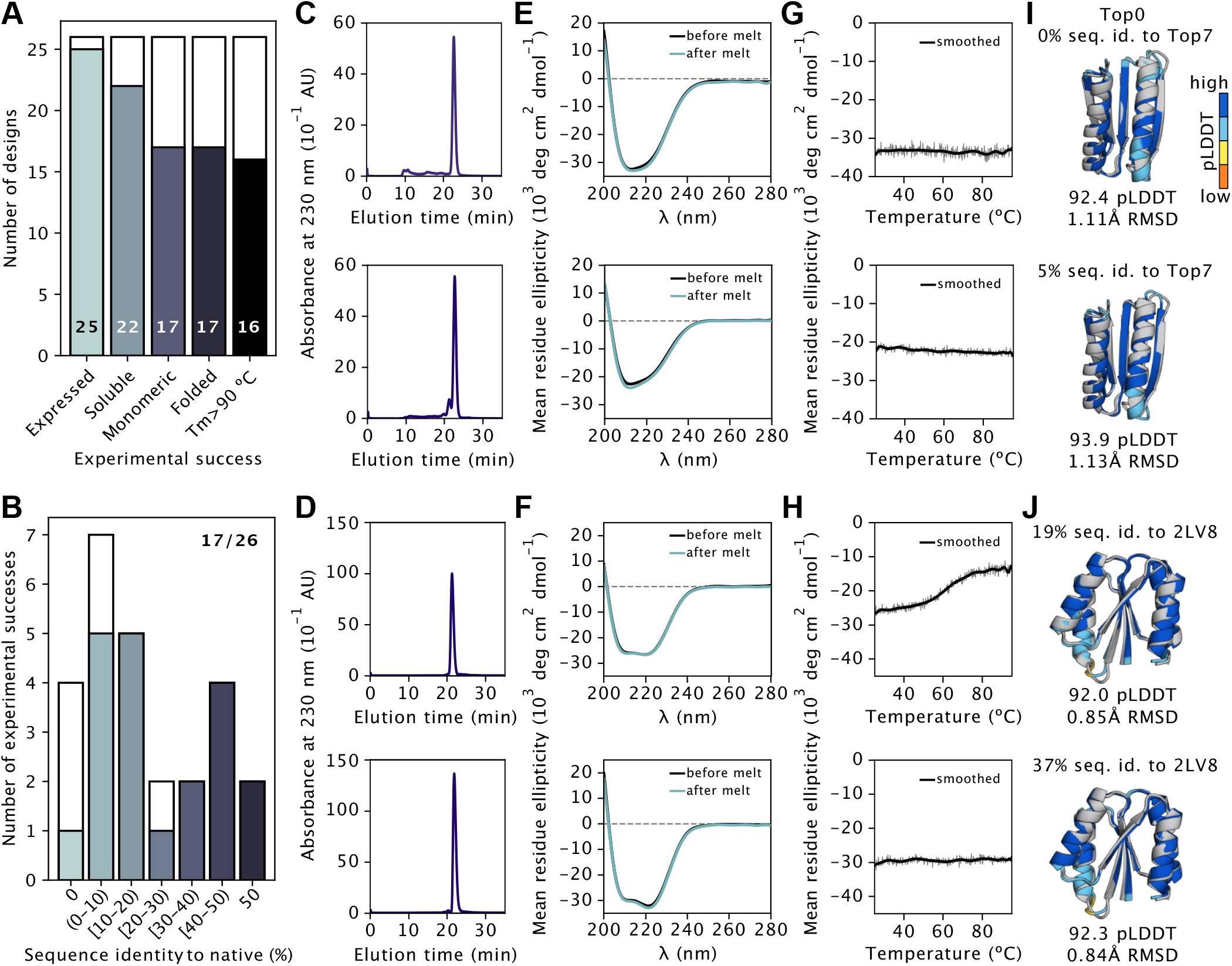
Experimental evaluation of Frame2seq designs. (A) Number of designs for the Top7 (PDBID 1QYS) and de novo Rossmann fold (PDBID 2LV8) backbones that achieve experimental success (filled bars). (B) Number of experimentally successful designs (filled bars) for different sequence identity bins. (C-H) Experimental characterization of four low sequence identity designs. (C) Size exclusion chromatography (SEC) profile for a 0% sequence identity (top) and 5% sequence identity (bottom) Top7 design. (D) SEC profile for a 19% sequence identity (top) and 37% sequence identity (bottom) 2LV8 design. (E) Circular dichroism (CD) spectra of a 0% (top) and 5% (bottom) Top7 design. (F) CD spectra of a 19% (top) and 37% (bottom) 2LV8 design. (G) Changes in CD signal (mean residue ellipticity) of a 0% (top) and 5% (bottom) Top7 design as a function of temperature. (H) Changes in CD signal of a 19% (top) and 37% (bottom) 2LV8 design as a function of temperature. (I) AlphaFold2 prediction for a 0% (top) and 5% (bottom) Top7 design (colored by pLDDT) aligned to the 1QYS X-ray structure (gray). (J) AlphaFold2 prediction for a 19% (top) and 37% (bottom) 2LV8 design (colored by pLDDT) aligned to the 2LV8 NMR structure (gray).

**Fig. 3.**
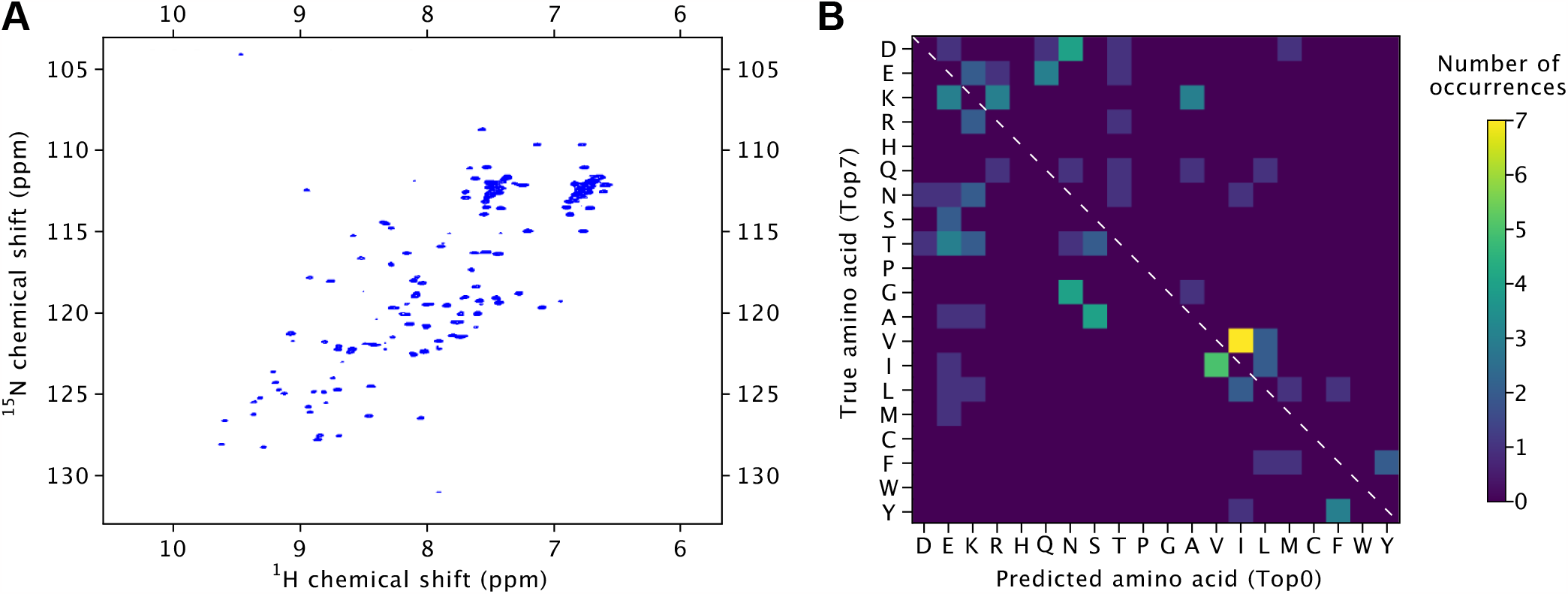
(A) The 2D [^15^N, ^1^H]-HSQC spectrum of 80µM ^15^N-labelled Top0 at pH = 6.7 recorded at 298.1K. (B) The substitution matrix computed position-wise between the true (Top7) and the predicted (Top0) amino acids. Number of occurrences is the total count of an amino acid substitution between the sequences. The white dashed line at the diagonal represents match of amino acid type.

## 4. Discussion

Protein design has greatly benefitted from advances in machine learning. Protein sequence design methods, such as ProteinMPNN (12) and PiFold (13), have reached unprecedented accuracy in terms of native sequence recovery. However, the state-of-the-art experimentally verified method, ProteinMPNN, has a slower inference speed than PiFold due to its autoregressive formulation. In this work, we present Frame2seq, a structure-conditioned masked language model for protein sequence design. We show that Frame2seq demonstrates improved accuracy with significantly reduced inference time compared to ProteinMPNN. Further, Frame2seq accurately estimates the error in its own predictions across diverse backbones, and model scores indicate awareness of “native-ness” during generation.

While native sequence recovery is a common measure of performance for protein design methods, the true measure of success for such a method would assess its ability to expand over the conditional sequence space exploited by nature without being limited to a single observed sequence. We assessed Frame2seq’s true success by designing for low sequence identity, down to zero percent. We found Frame2seq designs to be experimentally successful across all low identity bins. Our Top0 design, with zero percent sequence identity to Top7, displays biophysical characteristics consistent with a fold identical to Top7. Given backbones as input, Frame2seq successfully designs stable, soluble, and monomeric proteins that expand beyond the naturally occurring sequence space. We believe our method establishes a foundation for striking a balance between sufficient recovery of evolutionarily relevant sequence and sufficient divergence from a single observed example.

Exploration of novel sequences has become increasingly desirable with the rise of new de novo backbone generation methods, such as Chroma (8) and RFdiffusion (9). A common in silico pipeline for backbone generation methods is sequence design followed by structure prediction to assess structure self-consistency and designability. We believe Frame2seq’s speed and demonstrated novelty in sequence design will improve the success rates of these backbone generation methods and aid in the selection of designs for further experimentation.

A direct application of sampling maximally divergent sequences is for multi-state design objectives. Common techniques for multi-state design include reengineering of existing systems and directed evolution of proteins that do not generalize well or molecular dynamics simulations that are computationally costly. In addition, these techniques are limited by their inefficiency to output significant jumps in sequence space. Frame2seq coupled with structure prediction methods, such as AlphaFold2 (1), holds great promise in sampling from a jointly optimized sequence space where multiple folds share an identical sequence as in the case of fold-switching proteins. Frame2seq will be uniquely useful for multi-objective design applications where satisfaction of multiple functional criteria might require exploring larger sequence spaces to generate successful pareto-optimal solutions.

## Code and Data Availability

Frame2seq code is available at https://github.com/dakpinaroglu/Frame2seq

## Acknowledgments and Disclosure of Funding

We thank Jeffrey Ruffolo for insightful discussions and Dominic Grisingher for help setting up experiments. D.A. is funded by a National Science Foundation Graduate Research Fellowship (NSF GRFP). A.G. is a UCSF Discovery Fellow. T.K. is a Chan Zuckerberg Investigator.

## Supplementary Material

### A. Model details

#### A.1. Model architecture and training

The IPA components are based on the AlphaFold2 Structure Module PyTorch implementation from OpenFold (18). Our model contains 8 IPA layers followed by Structure Module transition layers. Unless it is the last IPA layer, the node transitions are followed by edge transitions (7 total) as implemented in (10). We set the hidden dimension to 128 and the Structure Module transition dimension to 128 (model 1) or 512 (models 2 and 3). Frame2seq models contain approximately 3.4M (model 1) or 10M (models 2 and 3) trainable parameters. Frame2seq models are trained with early stopping, for 144 or 200 epochs over the full dataset, which takes approximately 4 or 6 days on a single NVIDIA A40 GPU. We trained models using the Adam optimizer with the Noam scheduler.

**Table S1.** Frame2seq hyperparameters.

| Parameter | Value | Description |
| --- | --- | --- |
| $d_{\text{node}}$ | 128 | Node dimension |
| $d_{\text{edge}}$ | 128 | Edge dimension |
| $n_{\text{ipa-layers}}$ | 8 | IPA layers |
| $n_{\text{ipa-heads}}$ | 4 | IPA attention heads |
| $d_{\text{ipa-scalar-key}}$ | 16 | IPA scalar key dimension |
| $d_{\text{ipa-scalar-value}}$ | 16 | IPA scalar value dimension |
| $d_{\text{ipa-point-key}}$ | 4 | IPA point key dimension |
| $d_{\text{ipa-point-value}}$ | 6 | IPA point value dimension |

The IPA layers encode the sequence embedding, the edge embedding, and the coordinate frames, then predict and update the sequence embedding. The edge embeddings are updated by standard message passing with 2 fully connected layers.

We train models 1 and 2 with partial native sequence as input node features according to the following mask rate:

- With 75% probability, all amino acids are replaced with a masked residue token.
- With 25% probability, a sampled 0-100% of amino acids are provided for featurization.

#### A.2. Regularized invariant point attention layers

We adopt the invariant point attention from (1) and implement attention regularization via a dropout. Attention between pairs of residues is randomly masked at a 20% rate during training via the dropout in Algorithm 1 line 9.

##### Algorithm 1

Regularized invariant point attention (IPA)

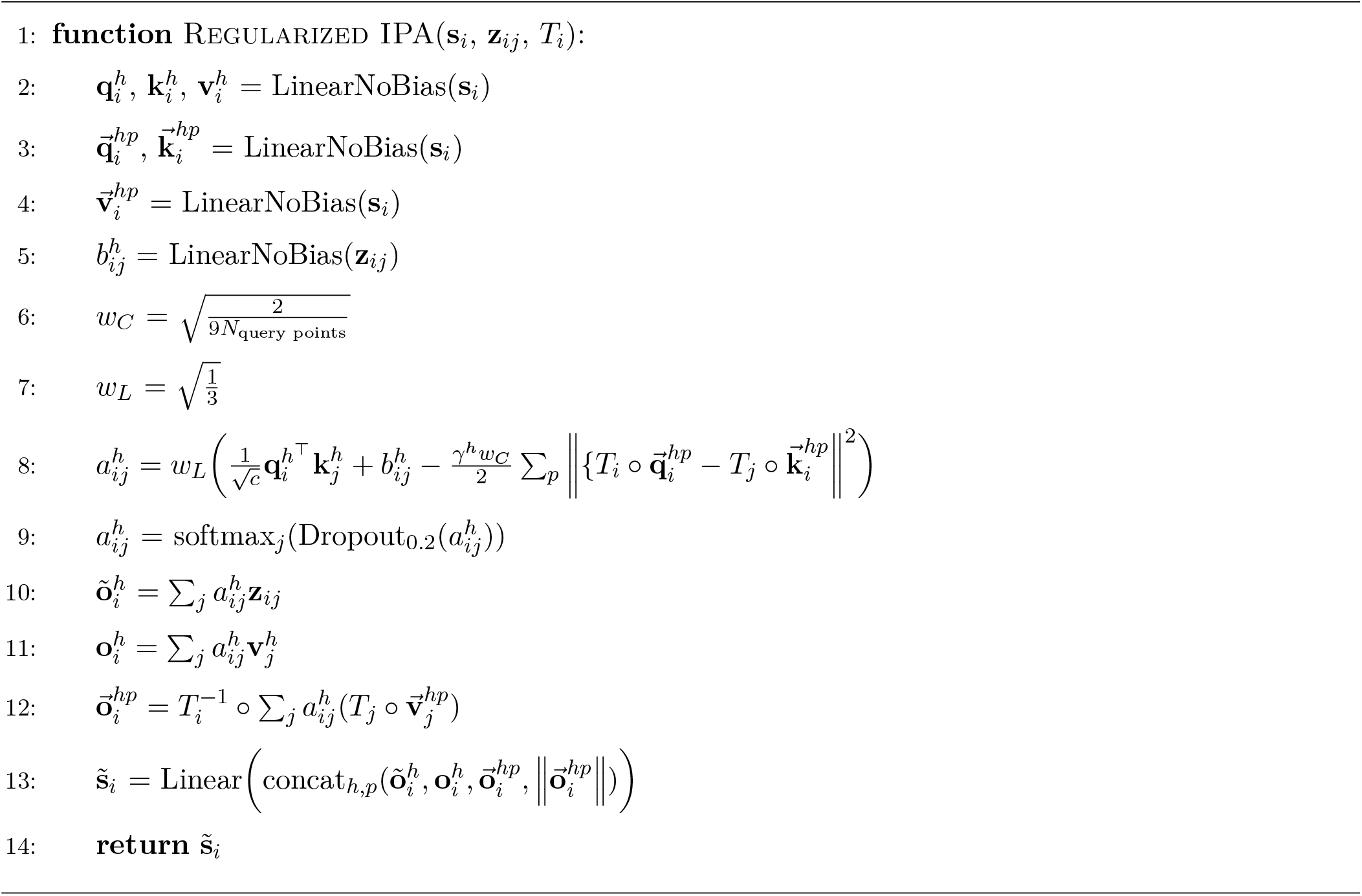

#### A.3. Ablation study

We ablate IPA regularization and edge update operations to study their effects on the performance of Frame2seq.

**Table S2.** Ablation study. Sequence recovery is median of average.

| Ablation | Sequence recovery<br>at 100 epochs (%) ↑ | Sequence recovery<br>at final epoch (%) ↑ |
| --- | --- | --- |
| No IPA regularization | 44.31 | — |
| No edge update operations | 44.32 | — |
| Baseline | 46.53 | — |
| Ensemble | 47.79 | **49.11** |

We ensemble 3 baseline models that are independently trained. We report Frame2seq’s performance as an ensemble.

### B. Further in silico analysis

The UMAP representations were obtained using n_neighbors = 100, metric =’euclidean’, min_dist = 0.3 and otherwise default parameters (Figure S1).

We used ColabFold (19) for all AlphaFold2 (1) predictions reported in this work (Figure S3).

**Fig. S1.**
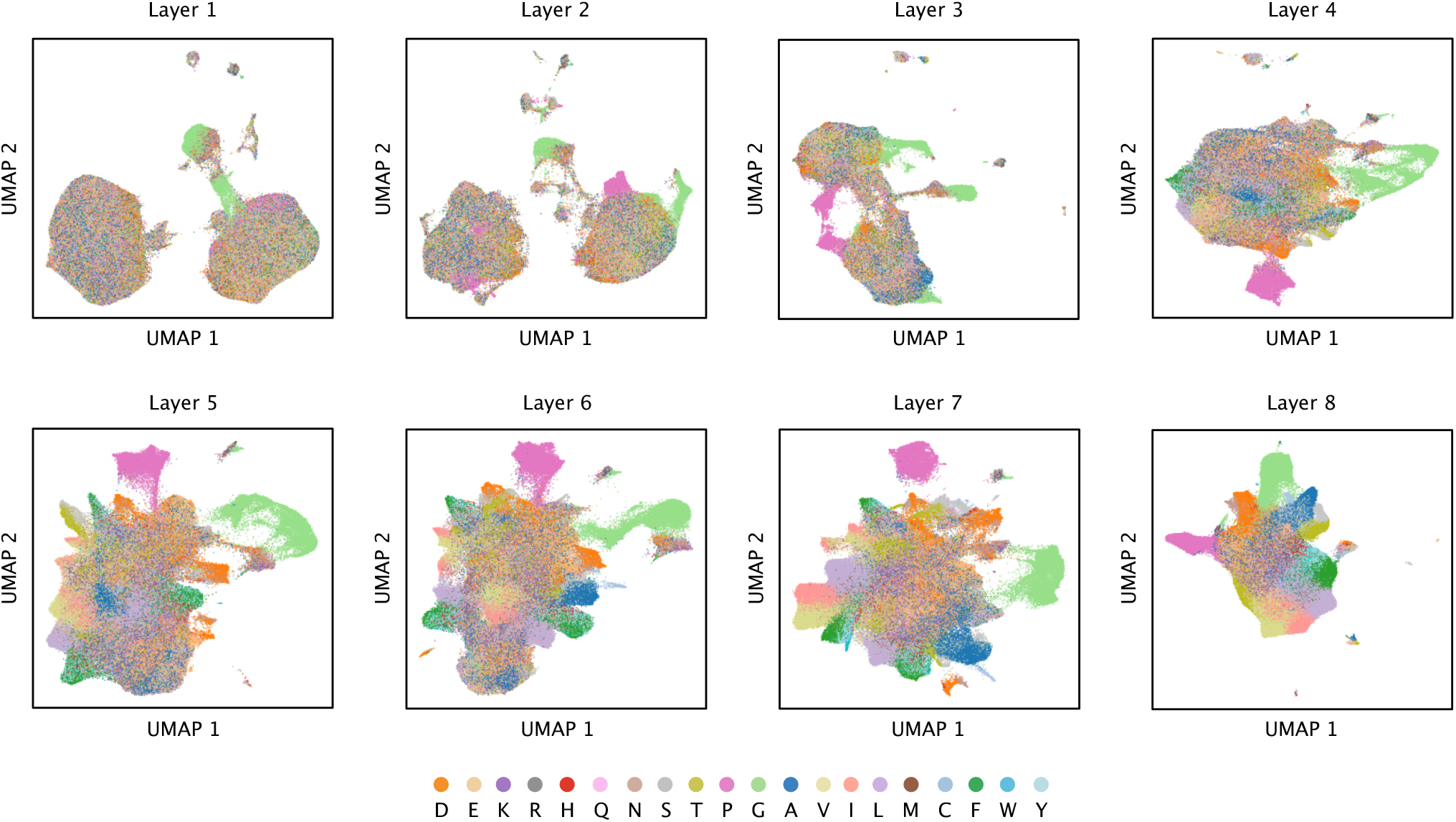
UMAP representation of Frame2seq embeddings from layers 1 through 8 for test set targets (colored by predicted amino acid identity).

**Fig. S2.**
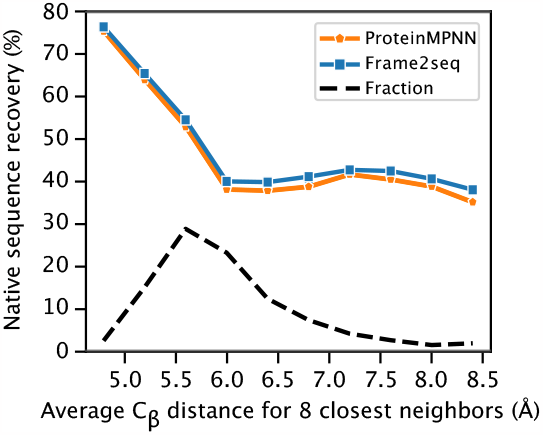
Native sequence recovery as a function of average C*β* distance for 8 closest neighbors.

**Table S3.** **Computational benchmark of model speed over CATH 4.2 held out test dataset targets. CPU speed (s) and GPU speed (s) is reported as mean of average for ProteinMPNN and Frame2seq inference**.

| Method | CPU speed (s) ↓ | GPU speed (s) ↓ |
| --- | --- | --- |
| ProteinMPNN | 13.70 | 2.71 |
| Frame2seq | **6.79** | **0.44** |

**Fig. S3.**
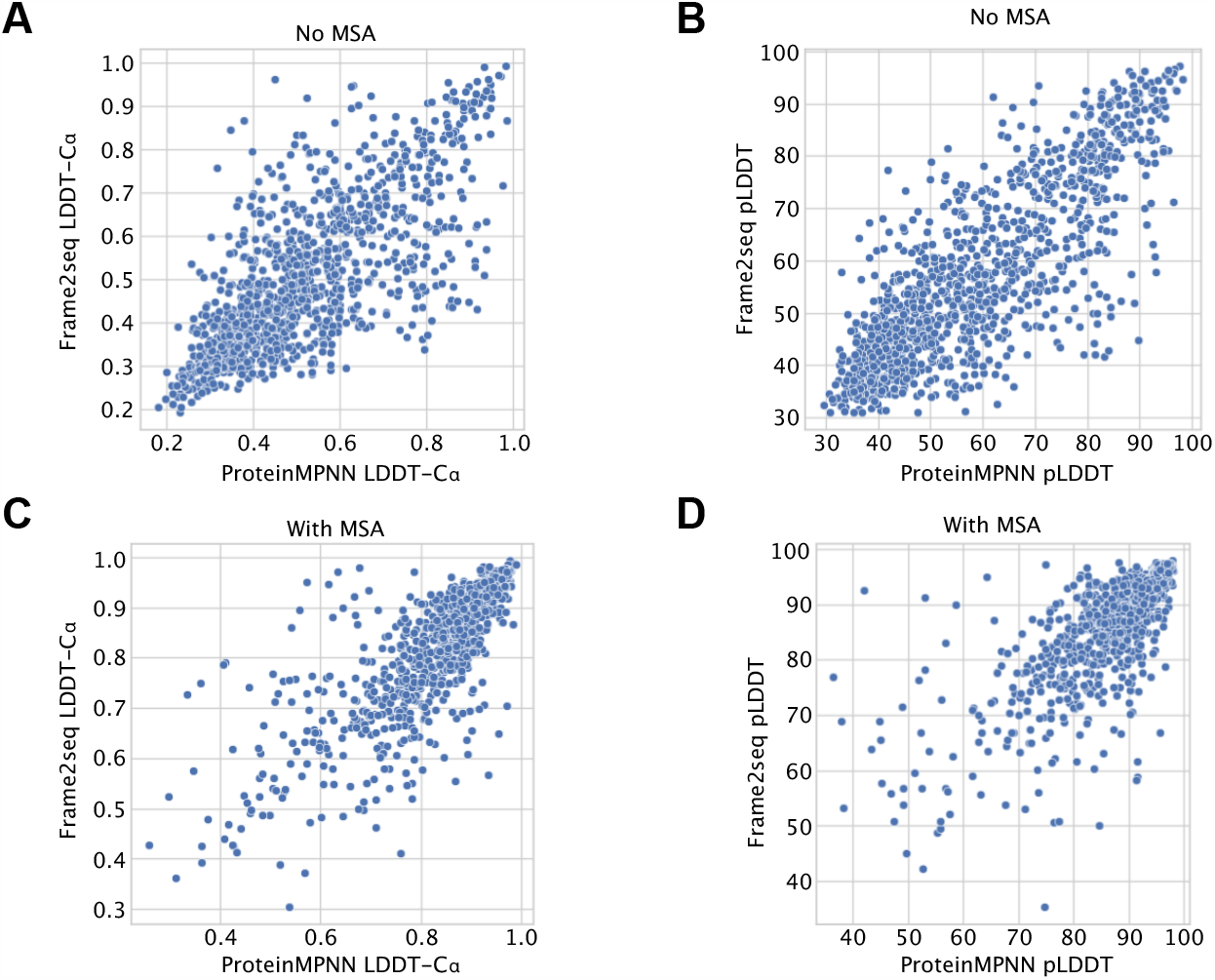
AlphaFold2 accuracy and confidence for ProteinMPNN and Frame2seq sequences. (A) AlphaFold2 accuracy (LDDT-C*α*) for predictions without MSAs (p-value = 0.052). (B) AlphaFold2 confidence (pLDDT) for predictions without MSAs (p-value = 0.012). (C) AlphaFold2 accuracy (LDDT-C*α*) for predictions with MSAs (p-value = 0.264). (D) AlphaFold2 confidence (pLDDT) for predictions with MSAs (p-value = 0.813).

**Fig. S4.**
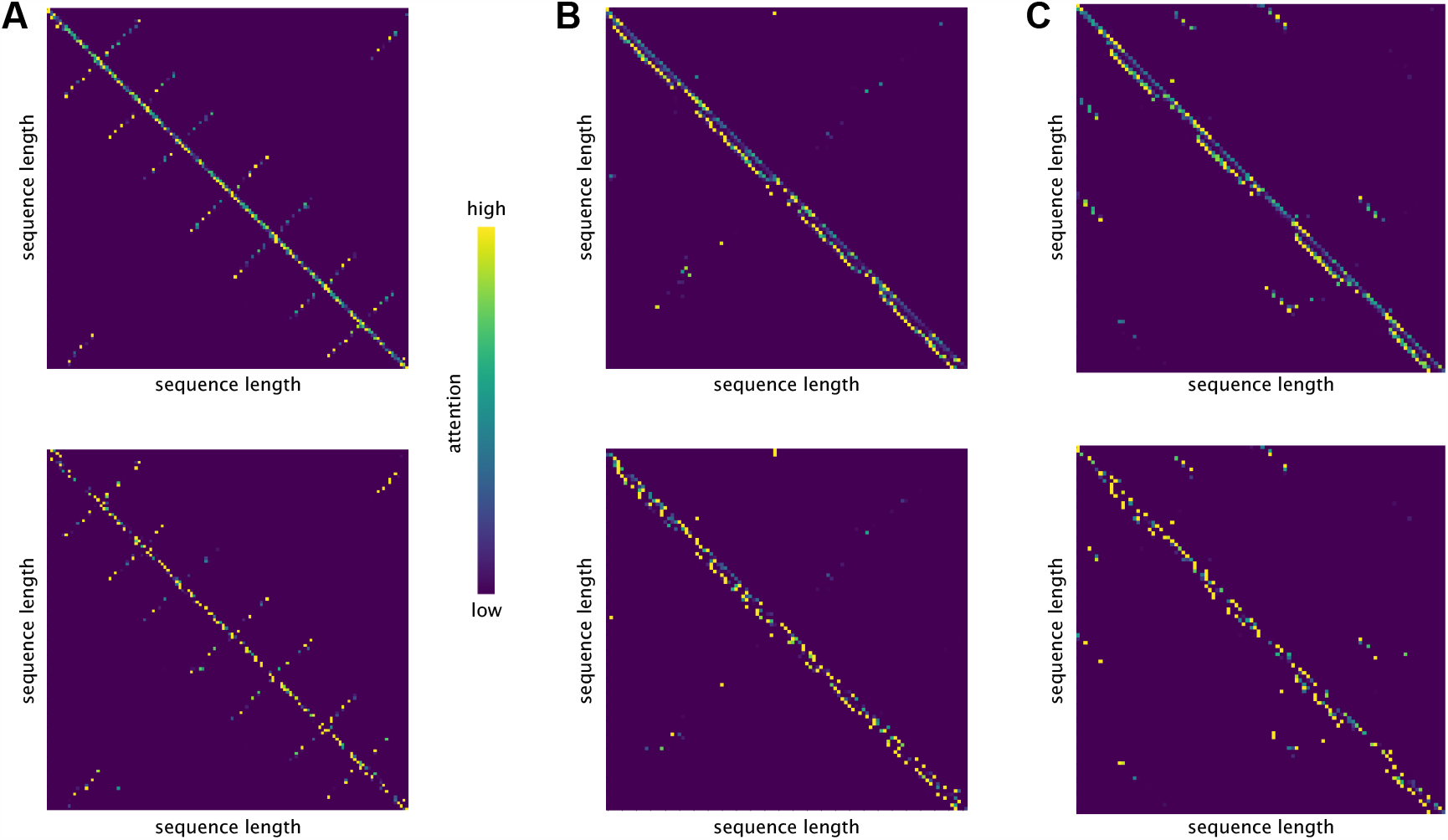
Effects of regularizing IPA on attention matrices. (A-C) Attention between pairs of residues with IPA (top) and with regularized IPA (bottom). (A) PDBID 6X1K. (B) PDBID 1P68. (C) PDBID 2LV8.

#### B.1. Design and selection of low sequence identity proteins

With true protein backbone as input, we used Frame2seq to sample designs with 0-50% sequence identity to the native. Cysteine was omitted from amino acid vocabulary during design. We filtered designs by total glutamic acid and aspartic acid count < total in native *±* 2. 10 out of 18 Top7 designs (set 2) were further filtered by a salt bridge count (matching native or higher in the helices) and a repulsive charge-charge interaction count (matching native or fewer in the helices).

**Fig. S5.**
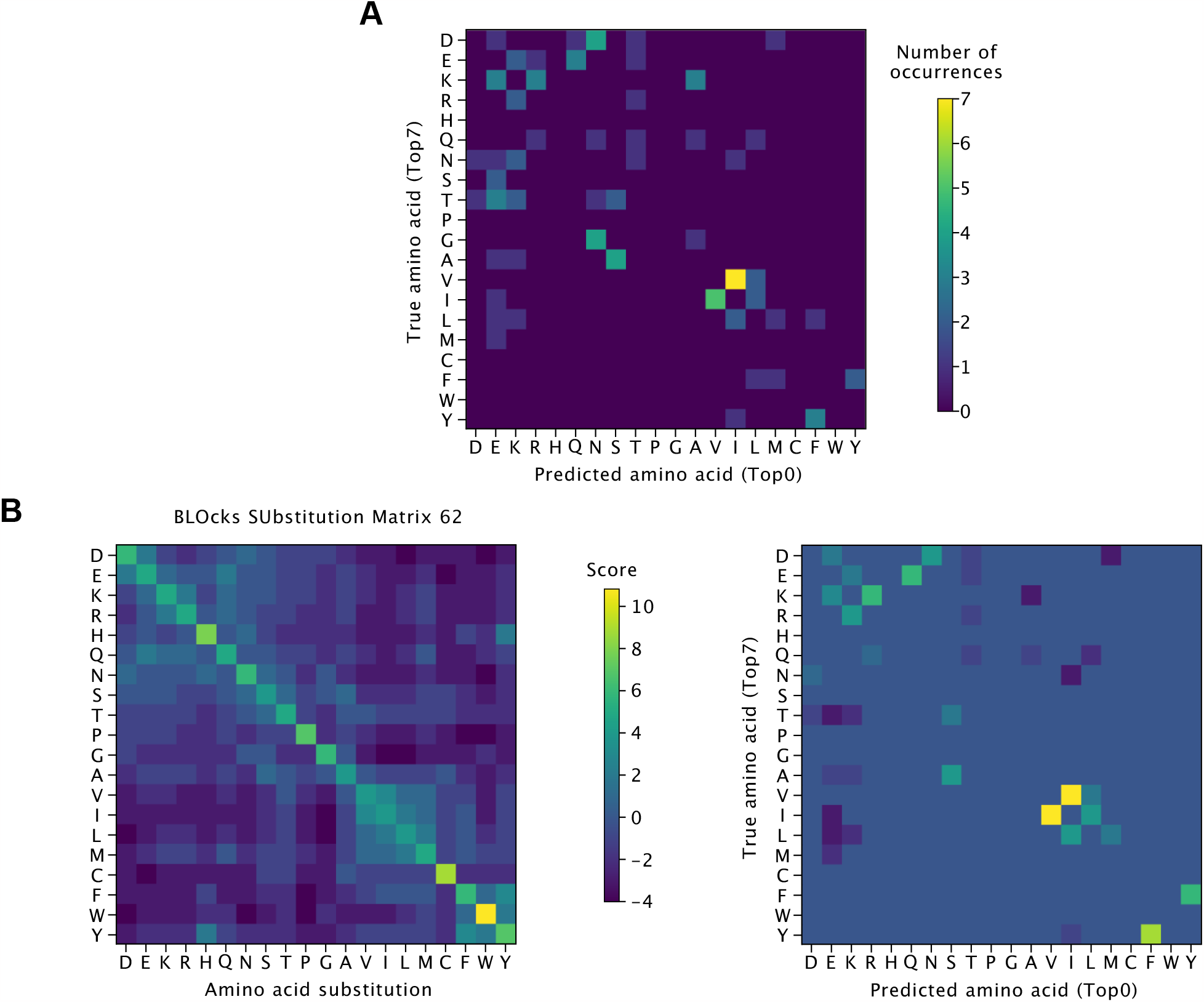
Comparison between Top7 and Top0 sequences. (A) The substitution matrix computed position-wise between the true (Top7) and the predicted (Top0) amino acids. Number of occurrences is the total count of an amino acid substitution between the sequences. (B) BLOcks SUbstitution Matrix (blosum) 62 for all possible amino acid substitutions (left) and blosum 62 score calculated between Top7 and Top0 (right).

### C. Experimental methods

#### C.1. Protein expression and purification

Designs were ordered with an N-terminal 6xHis-tag and thrombin cleavage site and were purchased from Twist Biosciences in a pET-28a+ vector. DNA was purified by transforming into DH5α *E. coli* cells, picking a single colony into LB-Kan[50], and miniprepping.

To screen for solubility and expression of the designs, chemically competent BL21(DE3) were transformed with DNA encoding designs using manufacturer’s protocol and plated onto LB-Kan[50] plates at 37 °C. A single colony was inoculated into 5 mL of LB-Kan[50] media at 37 °C or 18 °C with 220 rpm shaking overnight. The next day, 1 mL of culture was pelleted and stored at -80 °C, and protein expression was started in the remainder of the culture by adding 1 mM IPTG and growing at 30 °C or 18 °C. Cells were allowed to grow overnight, and were pelleted by centrifugation and stored at -80 °C. Cells were lysed either by using B-PER Bacterial Protein Extraction Reagent or by passing through a microfluidizer and centrifuged to pellet insoluble components. Lysate from uninduced cells, total induced cells, the supernatant of induced cells, and the pellet of induced cells were analyzed by SDS-PAGE to determine the solubility and expression level of protein designs.

For soluble designs, protein expression was scaled up. A single colony from a fresh transformation was inoculated into 5 mL of LB-Kan[50] media at 37 °C with 220 rpm shaking overnight. The next day, the entire overnight culture was added into either 250 mL or 500 mL of TB-Kan[50] with 220 rpm shaking at 37 °C. At OD = 0.4-0.6, protein expression was induced by adding IPTG to a final concentration of 1 mM. Temperature was reduced by transferring the cultures to a preheated 30 °C incubator, or by transferring cultures to a 4 °C cold room for 30 min before being transferred to a precooled 18 °C incubator. Cultures were allowed to grow overnight. Cells were harvested by centrifugation at 5,000 x g for 15 min at 4 °C.

Cell pellets were resuspended in 2 mL of resuspension buffer (30 mM sodium phosphate monobasic pH 7.5, 300 mM sodium chloride, 10 mM imidazole pH 7.5, 1 mg/mL Hen Egg White Lysozyme, and 15 U/mL Benzonase) per g of cell pellet. Cells were allowed to incubate in resuspension buffer for 30 minutes at room temperature with gentle shaking. Cells were additionally lysed either by passing the cell suspension through a microfluidizer three times, or by sonication on ice at 15% amplitude for 2 minutes using 5 s on / 5 s off cycles. Insoluble components were pelleted by centrifugation at 20 k x g for 20 min at 4 °C. Supernatant was decanted and incubated with 2 mL of pre-equilibrated 50% Ni-NTA resin slurry for 1 hr at 4 °C with gentle end-over-end shaking. The resin was washed four times with 20 mL of 30 mM sodium phosphate monobasic pH 7.5, 300 mM sodium chloride, and 20 mM imidazole pH 7.5. Proteins were eluted with 10 mL of 30 mM sodium phosphate pH 7.5, 300 mM sodium chloride, and 250 mM imidazole pH 7.5. Proteins were then dialyzed against at least 100X volumes of 1X PBS three times for at least 8 hours each at 4 °C, except for the 37% sequence identity design to 2LV8, which was dialyzed at room temperature. If large molecular weight contaminants were observed, they were removed using a 30 kDa MWCO Amicon concentrator and collecting the flow-through fraction. Protein concentrations were quantified by using a Bradford assay against a dilution series of a 1 mg/mL BSA standard or by measuring absorbance at 280 nm and calculating concentration using extinction coefficients as provided by ExPasy.

Protein purity was assessed by SDS-PAGE. Samples were prepared by mixing protein, 4X Laemmli buffer, 1 mM DTT, and water to a total volume of 15 µL and then denatured by incubation at 95 °C for 10 min. 10 µL of sample was loaded onto a 4-20% Tris-Glycine gel and run at 180 V for 45 min. Gels were stained with Coomassie dye, destained with water, and imaged using a Bio-Rad Gel-Doc EZ Imager.

#### C.2. Size-exclusion chromatography

Chromatography was done using an Agilent 1200 series HPLC attached to a Superdex S200 10/300 GL column and a UV detector. The UV detector was configured to detect signal at 230 nm using 360 nm as a baseline. 0.1 mg or 0.05 mg of protein with a concentration ranging from 0.2 to 3 mg/mL after purification was injected at a flow rate of 0.8 mL/min. All runs were done isocratically using 1X PBS as running buffer. Sizes were estimated using a linear regression model between elution time and molecular weight using BioRad Gel Filtration Standards.

#### C.3. Circular dichroism

Protein samples were diluted to 0.03 - 0.075 mg/mL in 1X PBS and added to a cuvette with a 1 or 2 mm path length. CD was completed using a Jasco J-710 spectrometer measuring from between 200 – 280 nm at a rate of 50 nm / min. Protein concentrations were adjusted as needed to keep the High-Tension Voltage within the linear range of the instrument. A thermal melt was performed from 25 – 95 °C at a ramp rate of 1 °C / min while measuring signal at 220 nm. Proteins were cooled back to 20 °C and were re-scanned to measure secondary structure after the thermal melt.

#### C.4. Protein expression for NMR structure determination

Double-labeled sample for NMR experiments was obtained by expressing in M9 minimal medium containing 4g/L D-glucose (U^13^C, 99%) and 0.5g/L 15NH4Cl (U^15^N, 99%). A single bacterial colony was inoculated into a 10mL starter culture grown at 37 °C overnight. The starter culture was diluted 1:50 into 1L ^13^C,-/^15^N-enriched M9 minimal medium and grown at 37 °C until OD_600_ reached 0.6-0.8. IPTG was then added to a final concentration of 300 µM to induce expression at 37 °C overnight. The expressed proteins were purified following the Ni NTA resin pull down protocol described in the protein purification section.

#### C.5. Acquisition of 2D 15N-HSQC spectrum

^13^C,-/^15^N-labeled protein was buffer exchanged into 25mM phosphate buffer (pH 6.7) and concentrated to 80µM before adding 5% (v/v) D_2_O. A two-dimensional ^15^N-HSQC (pulse program: fhsqcf3gpph) spectrum was recorded using a Bruker Avance NEO 800 MHz spectrometer with a 5mm TCI H&F-C/N-D CryoProbe at 298.1K. NMR spectra were processed using TopSpin 3.6.3 with indirect referencing to an external DSS standard (20).

**Fig. S6.**
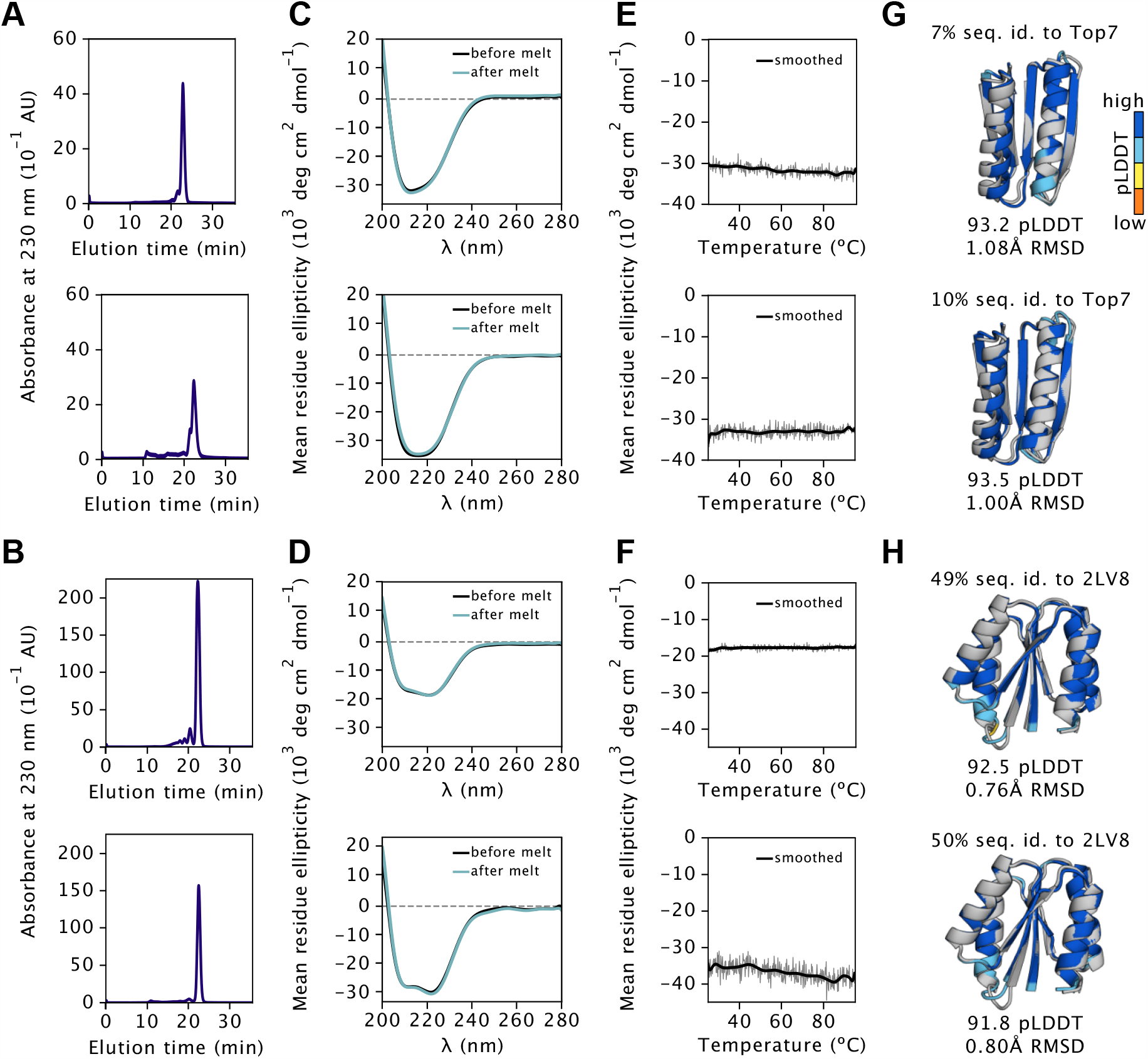
Further experimental evaluation of Frame2seq designs. (A) SEC profile for a 7% sequence identity (top) and 10% sequence identity (bottom) Top7 design. (B) SEC profile for a 49% sequence identity (top) and 50% sequence identity (bottom) 2LV8 design. (C) CD spectra of a 7% (top) and 10% (bottom) Top7 design. (D) CD spectra of a 49% (top) and 50% (bottom) 2LV8 design. (E) Changes in CD signal of a 7% (top) and 10% (bottom) Top7 design as a function of temperature. (F) Changes in CD signal of a 49% (top) and 50% (bottom) 2LV8 design as a function of temperature. (G) AlphaFold2 prediction for a 7% (top) and 10% (bottom) Top7 design (colored by pLDDT) aligned to the 1QYS X-ray structure (gray). (H) AlphaFold2 prediction for a 49% (top) and 50% (bottom) 2LV8 design (colored by pLDDT) aligned to the 2LV8 NMR structure (gray).

**Table S4.**
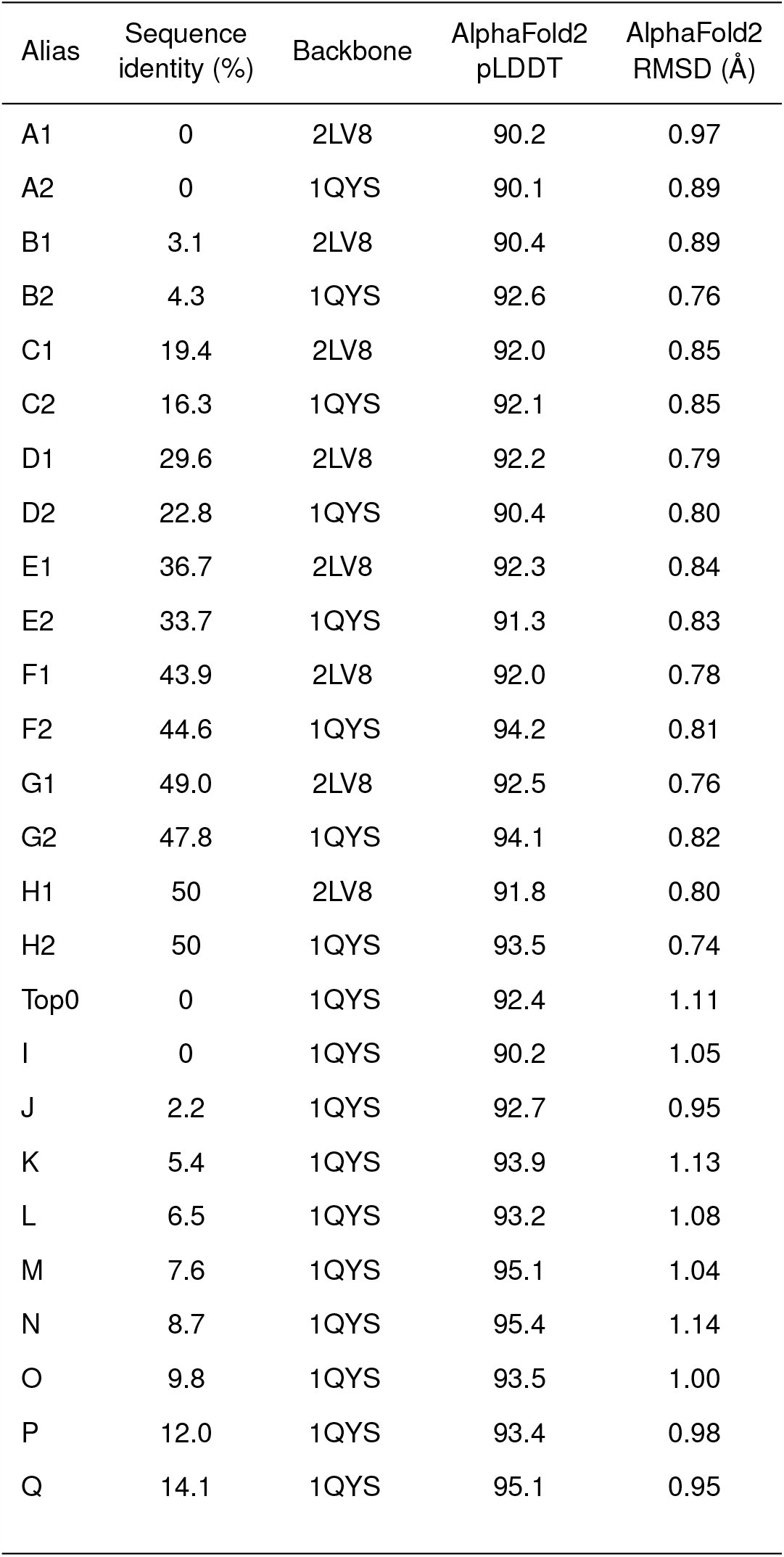
Sequence identity to native, AlphaFold2 pLDDT, and AlphaFold2 RMSD to native for low sequence identity designs.

| Alias | Sequence identity (%) | Backbone | AlphaFold2 pLDDT | AlphaFold2 RMSD (Å) |
| --- | --- | --- | --- | --- |
| A1 | 0 | 2LV8 | 90.2 | 0.97 |
| A2 | 0 | 1QYS | 90.1 | 0.89 |
| B1 | 3.1 | 2LV8 | 90.4 | 0.89 |
| B2 | 4.3 | 1QYS | 92.6 | 0.76 |
| C1 | 19.4 | 2LV8 | 92.0 | 0.85 |
| C2 | 16.3 | 1QYS | 92.1 | 0.85 |
| D1 | 29.6 | 2LV8 | 92.2 | 0.79 |
| D2 | 22.8 | 1QYS | 90.4 | 0.80 |
| E1 | 36.7 | 2LV8 | 92.3 | 0.84 |
| E2 | 33.7 | 1QYS | 91.3 | 0.83 |
| F1 | 43.9 | 2LV8 | 92.0 | 0.78 |
| F2 | 44.6 | 1QYS | 94.2 | 0.81 |
| G1 | 49.0 | 2LV8 | 92.5 | 0.76 |
| G2 | 47.8 | 1QYS | 94.1 | 0.82 |
| H1 | 50 | 2LV8 | 91.8 | 0.80 |
| H2 | 50 | 1QYS | 93.5 | 0.74 |
| Top0 | 0 | 1QYS | 92.4 | 1.11 |
| I | 0 | 1QYS | 90.2 | 1.05 |
| J | 2.2 | 1QYS | 92.7 | 0.95 |
| K | 5.4 | 1QYS | 93.9 | 1.13 |
| L | 6.5 | 1QYS | 93.2 | 1.08 |
| M | 7.6 | 1QYS | 95.1 | 1.04 |
| N | 8.7 | 1QYS | 95.4 | 1.14 |
| O | 9.8 | 1QYS | 93.5 | 1.00 |
| P | 12.0 | 1QYS | 93.4 | 0.98 |
| Q | 14.1 | 1QYS | 95.1 | 0.95 |

**Table S5.**
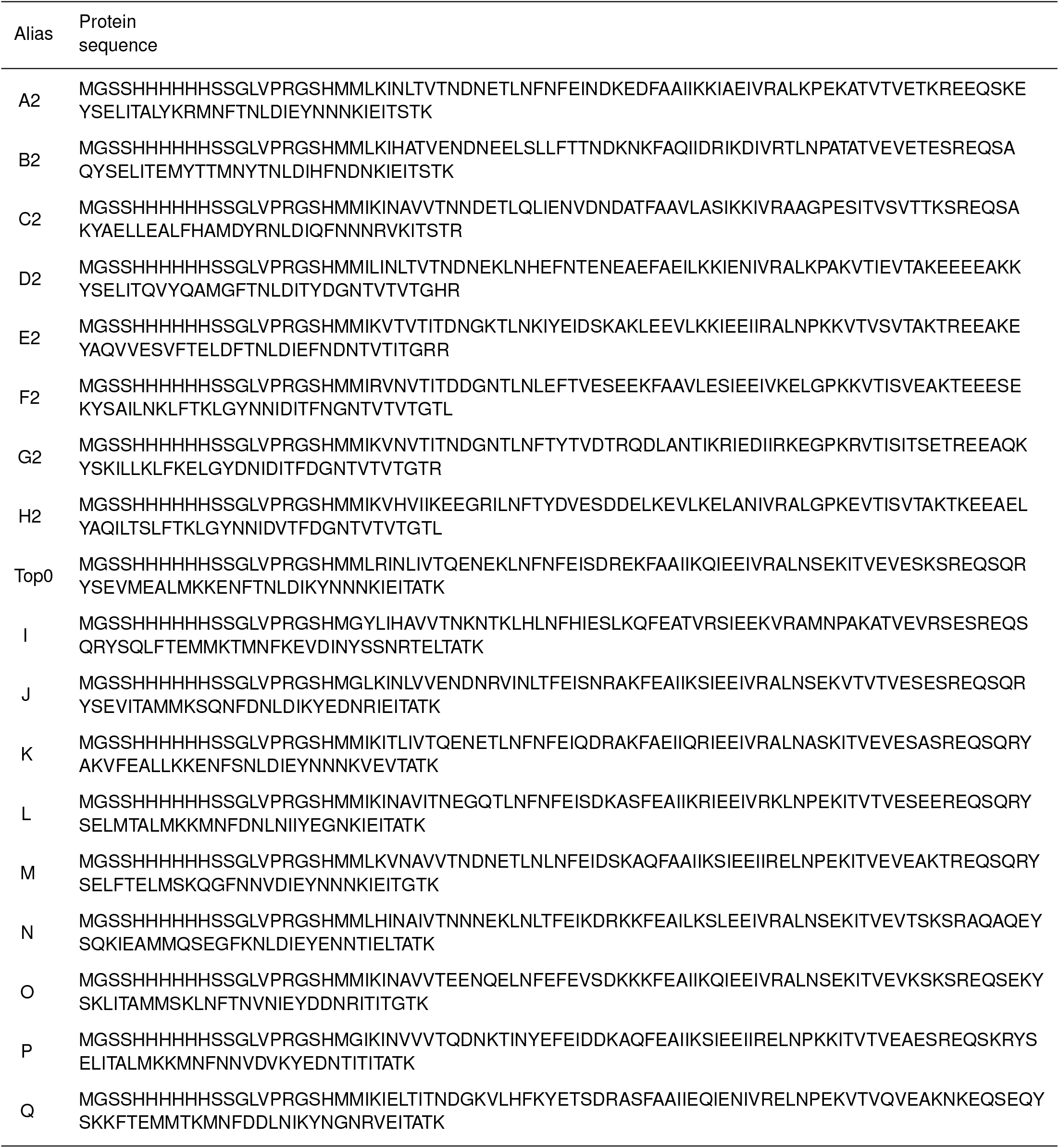
Protein sequences for low sequence identity 1QYS designs.

**Table S6.**
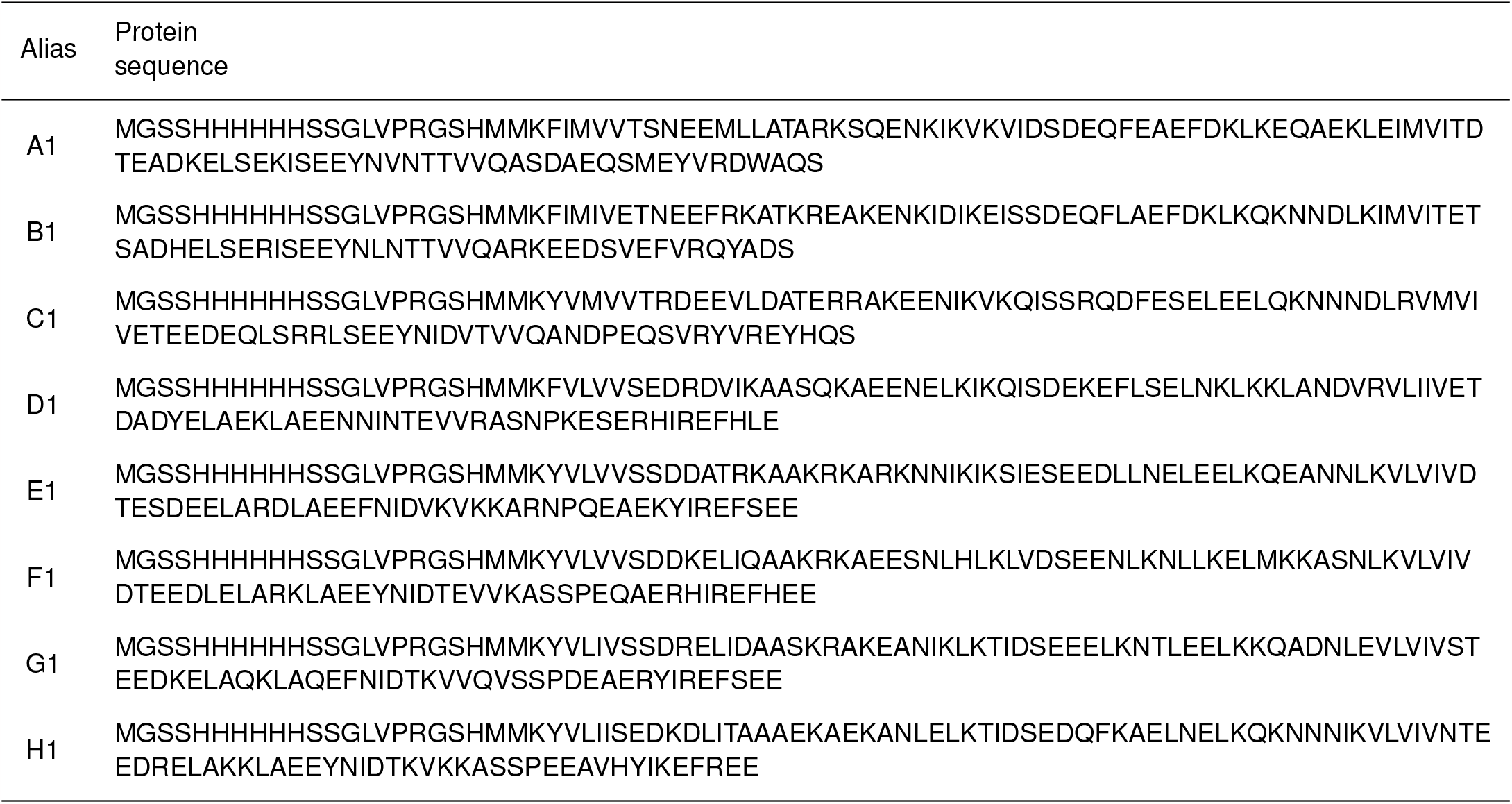
Protein sequences for low sequence identity 2LV8 designs.

**Table S7.**
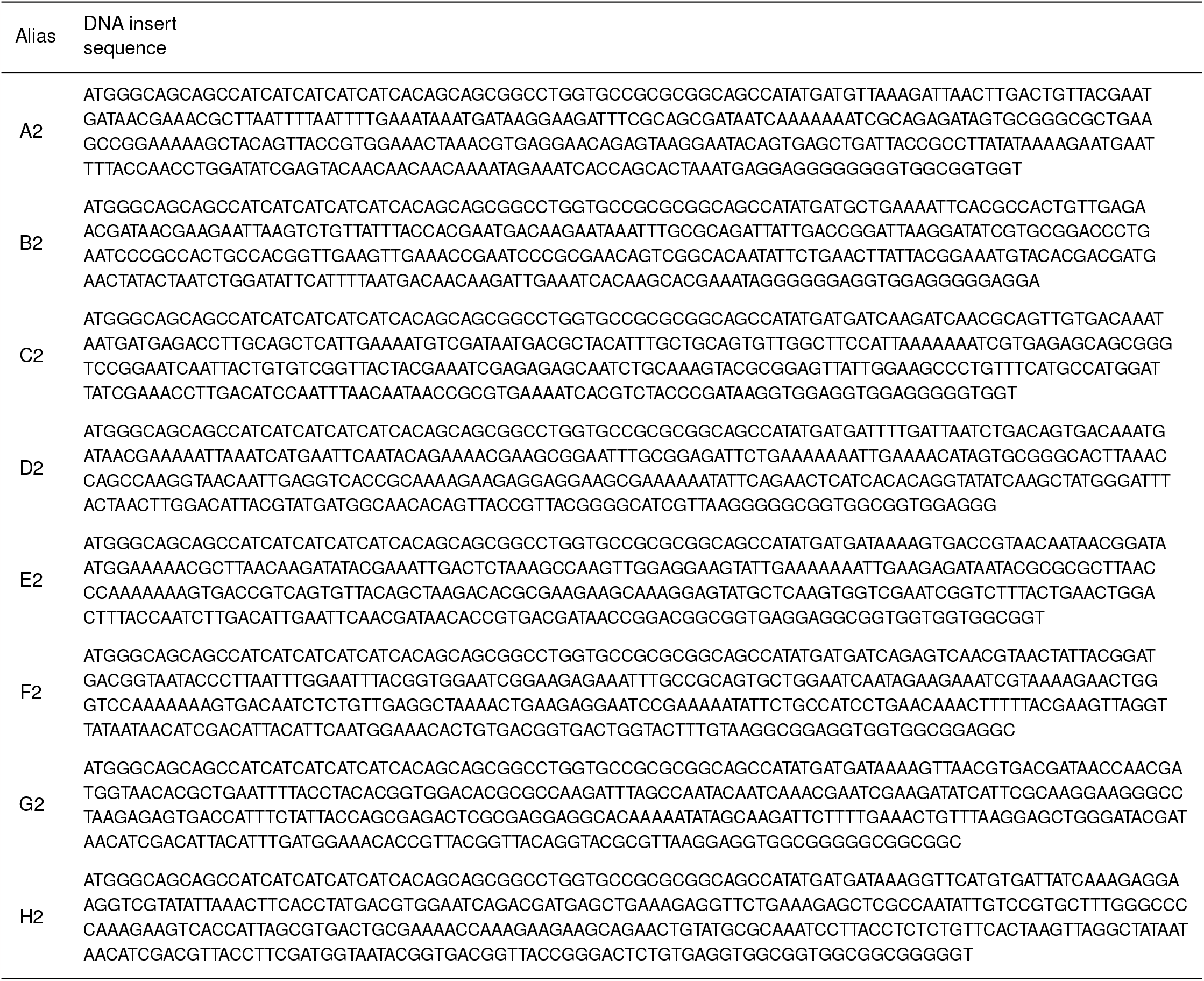
DNA insert sequences for low sequence identity 1QYS designs in set 1.

**Table S8.**
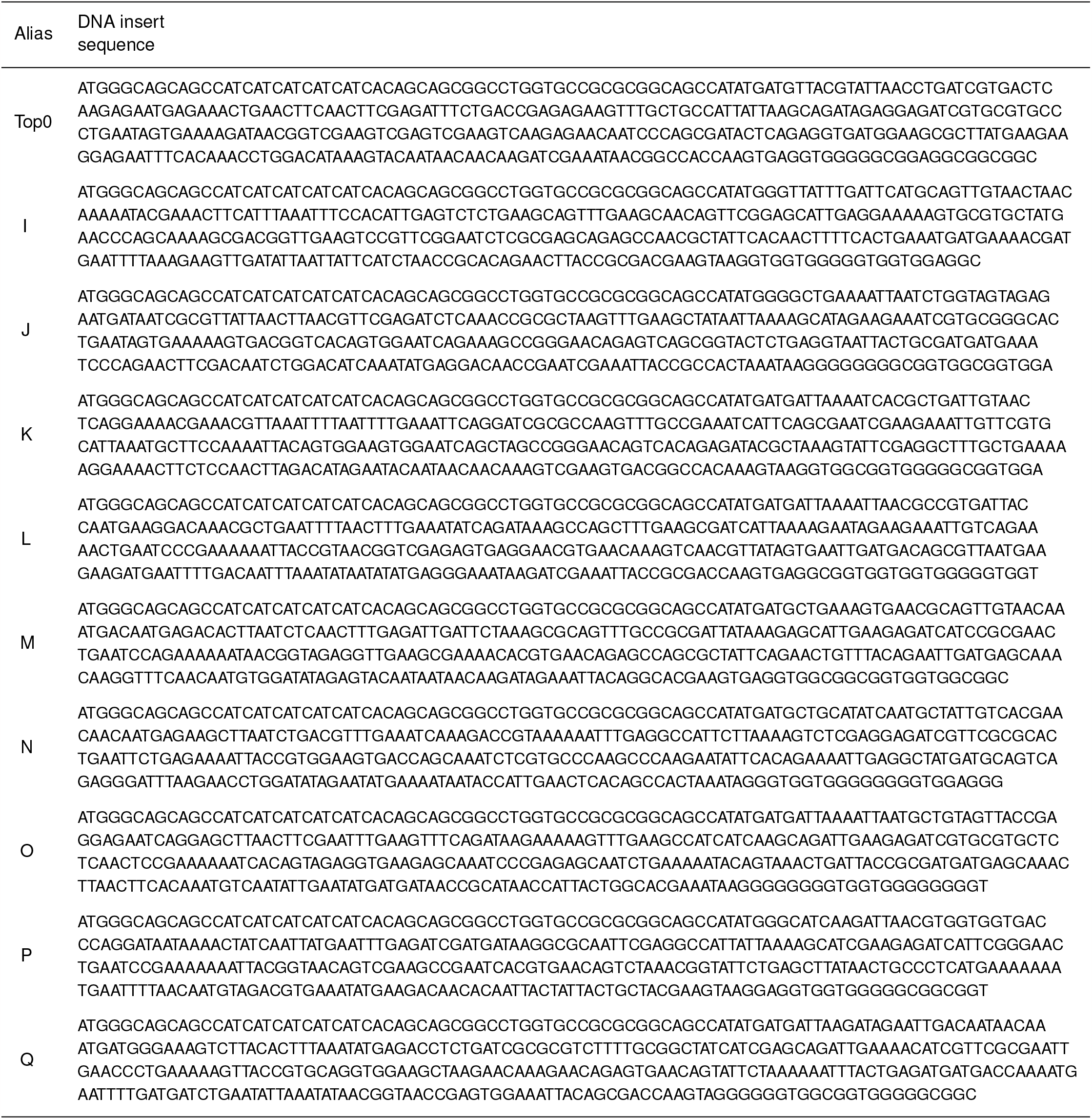
DNA insert sequences for low sequence identity 1QYS designs in set 2.

**Table S9.**
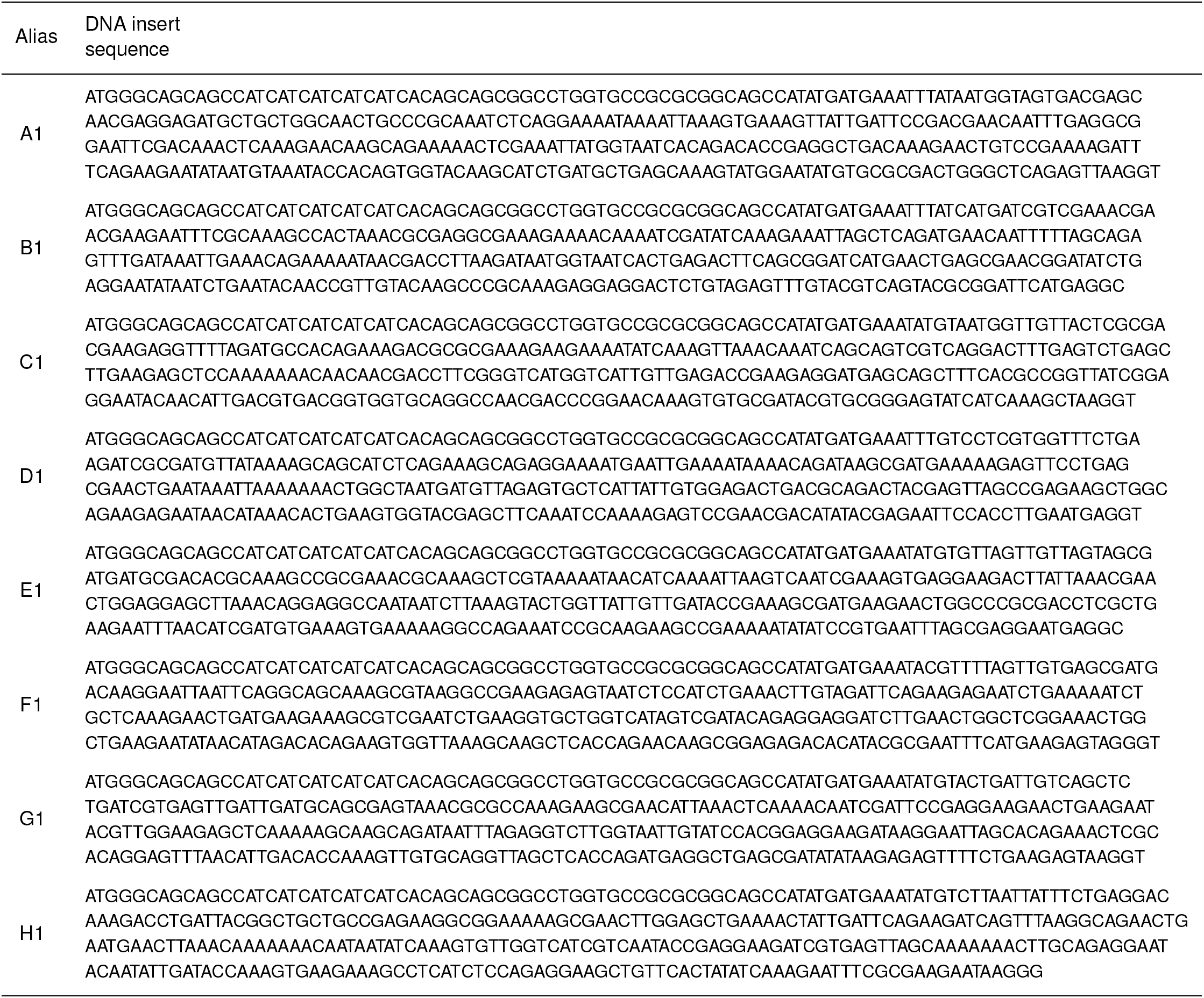
DNA insert sequences for low sequence identity 2LV8 designs.

